# Direct single-molecule visualization of Hsp90-mediated relief of a Hsp70-folding block

**DOI:** 10.1101/2025.08.25.672245

**Authors:** Nicholas R. Marzano, Bailey Skewes, Shannon McMahon, Lauren Rice, Dezerae Cox, Antoine M. van Oijen, Heath Ecroyd

## Abstract

Hsp70 and Hsp90 are ubiquitous molecular chaperones that cooperate to promote the correct folding and maturation of client proteins. Despite their central role in proteostasis, the molecular mechanisms by which Hsp90 coordinates with Hsp70 to remodel clients remain poorly understood. In particular, how ATP hydrolysis by Hsp90 is coupled to client engagement and conformational change has been a long-standing question. This gap in understanding is largely due to the challenge of visualizing client conformations during the highly dynamic and heterogeneous interactions with the Hsp70/Hsp90 chaperone machinery. To address this, we used a combination of single-molecule fluorescence resonance energy transfer (smFRET) and total internal reflection fluorescence microscopy to observe individual firefly luciferase client proteins as they are sequentially engaged by Hsp70 and Hsp90. Here, we show that Hsp90 reduces rebinding of Hsp70 to folding intermediates while still allowing engagement to misfolded clients, thereby enabling productive refolding in the presence of typically inhibitory concentrations of Hsp70. Furthermore, Hsp90 uses ATP binding and hydrolysis to actively remodel the conformational landscape of the client, promoting controlled folding through localized compaction across multiple regions. These controlled folding events reduce misfolding and are essential for establishing native interdomain contacts. Using smFRET and kinetic simulations, we further demonstrate that heterogeneous Hsp70 binding generates region-specific folding kinetics and conformational dynamics, which are likely driven by variations in the number of available Hsp70-binding sites. This is then exploited during Hsp70/Hsp90-mediated folding to support localized folding of client subdomains, thereby reducing non-native interactions from distal regions to facilitate proper folding of multi-domain proteins.

## Introduction

Most proteins need to fold into specific, three-dimensional (native) conformations to perform their biological functions ^1^. In the cell, however, folding represents a challenging problem for a protein due to the large number of conformations that a polypeptide can adopt ^2^ and their propensity to enter ‘off-folding’ pathways, which can lead to the accumulation of non-native, misfolded structures. These can result in diseases associated with protein loss-of-function (*e.g.*, Cystic Fibrosis) ^3^ or protein aggregation, including numerous neurodegenerative disorders (*e.g.*, Alzheimer’s and Parkinson’s disease) ^4,5^. Consequently, cells have evolved a vast and diverse network of molecular chaperones to assist in the *de novo* folding and refolding of misfolded proteins to their native conformations ^6^. Among the most studied molecular chaperones are the intracellular heat shock proteins (Hsps), notably the Hsp70 and Hsp90 systems, which work cooperatively to guide polypeptides toward their native state ^7,8^.

The current model by which Hsp70 and Hsp90 cooperate to fold proteins is best established for the *Escherichia coli* (*E. coli*) system, whereby a Hsp40 co-chaperone (DnaJ) binds to a client protein and delivers it to ATP-bound Hsp70 (DnaK). DnaJ stimulates ATP hydrolysis 1000 fold ^9^, which causes large-scale conformational rearrangements in DnaK that result in ultra-stable capture of the client protein by Hsp70. Binding of multiple Hsp70 proteins to a single client induces substantial client expansion to resolve misfolded states ^10,11^. Association of a nucleotide-exchange factor (NEF, GrpE in *E. coli*) promotes exchange of ADP for ATP on Hsp70, which promotes release of the client and provides an opportunity for productive folding; multiple cycles of client binding-and-release eventually drive folding to the native state ^11^. However, recent evidence suggests that high, yet physiologically-relevant, concentrations of Hsp70 promote excessive binding of the client protein by Hsp70, which effectively blocks refolding; when this occurs, transfer of the client to Hsp90 (HtpG) alleviates Hsp70-induced inhibition and allows folding to proceed ^12^. It is well established that Hsp90 function is driven by large-scale conformational rearrangements associated with ATP binding and/or hydrolysis, with many structural and single-molecule analyses describing these conformational changes in Hsp90 ^13–15^. However, there are a number of outstanding questions that remain with regard to the client protein during cooperative folding by Hsp70 and Hsp90 including: (i) how do the nucleotide-regulated changes in the chaperones influence the conformation of the client protein so as to promote protein folding?; (ii) how does Hsp90 protect the client from Hsp70-mediated inactivation?; and (iii) what is the role of ATP binding *versus* hydrolysis on the conformation of the client protein?

A difficulty in addressing these outstanding questions is the heterogeneous, dynamic and transient nature by which these chaperones interact with client proteins, which substantially complicates efforts to identify and characterize important processes. Whilst NMR ^16–18^ and structural studies ^19–21^ have been used to visualize the conformation of clients bound to Hsp90, these only provide static representations of proteins trapped in defined states (e.g., ATP vs ADP bound). As such, real-time observation of client conformational dynamics throughout the entire Hsp70/Hsp90 cycle is not possible from protein structures alone and the contribution(s) of these dynamics to protein folding are not well understood. To overcome this limitation, we performed single-molecule fluorescence resonance energy transfer (smFRET) experiments to temporally monitor the conformation of individual molecules of the multi-domain client protein firefly luciferase (Fluc) as it is sequentially engaged and folded by the bacterial Hsp70 and Hsp90 systems.

We find that HtpG enables clients to escape DnaK-mediated folding inhibition by reducing DnaK client binding and promoting controlled, stepwise folding events that are driven by ATP hydrolysis, thereby enhancing the probability of correct folding and reducing misfolding. These mechanisms are conserved across different domains of Fluc, suggesting a general mechanism by which HtpG uses ATP hydrolysis to fold clients. smFRET and kinetic simulations indicate that DnaK binds dynamically to multiple sites on Fluc, with regional differences in the number of DnaK binding sites controlling client expansion kinetics; this enables DnaK and HtpG to spatially separate domains, allowing subdomains to fold independently and efficiently. Collectively, Hsp70/Hsp90-mediated folding provides a mechanism to rescue kinetically trapped proteins and subsequently smooth the folding landscape, thereby enabling rapid client folding with high fidelity.

## Methods

### Materials

Plasmids encoding N-terminally SUMO-tagged Fluc mutants with C-terminal AviTag were generated previously ^11^ or modified via site-directed mutagenesis by Genscript. Plasmids encoding chaperones were kindly donated by M. Mayer (University of Heidelberg, Germany).

### Protein expression and purification

Fluc isoforms were expressed and purified as previously described ^11^ with the following modification: upon initial purification of the cell lysate via immobilized metal affinity chromatography (IMAC), bound protein was first washed with 10 mM D-biotin for 1 h to dissociate non-specific Fluc-BirA complexes prior to elution with imidazole. DnaK, DnaJ and GrpE were also purified as described previously ^11^ without modification.

Unless otherwise specified, a modified form of HtpG was used that contains the coiled-coil motif of the kinesin neck-region of *Drosophila melanogaster* to the C-terminus of HtpG (to maintain a dimeric state) as well as a C-terminal Strep Tag II for purification ^13^. Briefly, BL21(DE3) cells containing the plasmid encoding HtpG were grown to mid-log phase (0.6 – 0.8 optical density at 600 nm) in 2x YT media (16 g/L tryptone, 10 g/L yeast and 5 g/L NaCl, pH 7.0) supplemented with ampicillin (100 µg/mL) at 37°C and protein expression induced by addition of and 0.1% (w/v) L-arabinose. Induced cultures were then returned to the orbital shaker for agitation (180 rpm) at 30°C overnight. Cells were subsequently harvested by centrifugation at 5000 × *g* for 20 min at 4°C, with the obtained cell pellets stored at −20°C until further purification took place. Cells were resuspended in lysis buffer (100 mM Tris-HCl [pH 8.0], 150 mM NaCl, 1 mM EDTA, 2mM β-mercaptoethanol [BME]) supplemented with cOmplete™ Protease Inhibitor Cocktail (Sigma) and lysozyme (0.5 mg/mL) and resuspended pellets were left rocking at 4°C for 30 min. DNase I was added to lysates at a final concentration of 5 mg/mL and incubated at 4°C for a further 30 min with rocking. The lysate was probe sonicated for 3 min (10s on/20s off) at 45% power and then clarified via two rounds of centrifugation at 24,000 × *g* at 4°C for 20 min. The soluble fraction was taken and filtered through a 0.45 µm and loaded onto a 5 mL StrepTrap XT Sepharose column (Cytiva) pre-equilibrated in lysis buffer. Unbound proteins were then washed from the column with lysis buffer and recombinant protein eluted following the addition of lysis buffer supplemented with 50 mM biotin. Fractions containing recombinant protein were pooled and dialyzed at 4°C overnight into storage buffer (100 mM Tris-HCl [pH 7.5], 150 mM NaCl, 1 mM EDTA). Dialyzed protein was then concentrated using a Pierce™ Protein Concentrator (10K MWCO) at 4,000 × *g* at 4°C, snap-frozen and stored at –80°C until required.

HtpG^WT^ (which does not contain the coiled-coil motif), HtpG^E34A^ (which has reduced ATP hydrolysis) and HtpG^R355L^ (DnaK-binding defective) were expressed in *ΔHtpG* cells (containing a deletion of wild-type HtpG) as described above, with the exception that cells were induced upon addition of 1 mM IPTG. Cells were resuspended in IMAC lysis buffer (40mM HEPES-KOH [pH 7.5], 100 mM KCl, 5 mM MgCl_2_, 10% (v/v) glycerol, 4 mM BME, 20 mM imidazole) supplemented with cOmplete™ Protease Inhibitor Cocktail (Sigma) and lysozyme (0.5 mg/mL) and soluble protein extracted as described above. The entire lysate (containing recombinant protein with N-terminal 6xHis-SUMO tag) was loaded onto a 5 mL His-Trap Sephadex column (Cytiva) pre-equilibrated in HtpG IMAC lysis buffer and washed in the same buffer. Bound recombinant protein was eluted with the addition of IMAC lysis buffer supplemented with 250 mM imidazole. Collected eluate was analyzed via SDS-PAGE and protein-containing fractions were pooled and dialyzed at 4°C overnight into cleavage buffer (40mM HEPES-KOH (pH 7.5), 50 mM KCl, 5 mM MgCl_2_, 10% (v/v) glycerol, 4 mM BME, 20 mM imidazole] with the addition of Ulp1 protease (final concentration: 4 µg/mg of recombinant protein) to remove the 6xHis-SUMO tag. Cleaved protein was then returned to the IMAC column and purified as described above, with fractions containing cleaved recombinant protein pooled and dialyzed into HtpG storage buffer (40 mM HEPES-KOH [pH 7.5], 150 mM KCl, 5 mM MgCl_2_, 10% (v/v) glycerol, 4 mM BME). Protein was then concentrated using a Pierce™ Protein Concentrator (10K MWCO) at 4,000 × *g* at 4°C, snap-frozen and stored at –80°C until required.

### Protein labeling

Fluc constructs were fluorescently labeled with an Alexa Fluor 555 (AF555) and Alexa Fluor 647 (AF647) FRET pair as described previously ^22^ with minor modifications. Briefly, Fluc (2 mg/ml) was incubated in the presence of 5 mM tris(2-carboxyethyl)phosphine and 40% (w/v) ammonium sulfate and placed on a rotator for 1 hour at 4°C. Proteins were then centrifuged at 20,000 x *g* for 15 min and resuspended in degassed buffer A (100 mM Na_2_PO_4_ [pH 7.4], 200 mM NaCl, 1 mM EDTA, and 40% [w/v] ammonium sulfate). The protein was then centrifuged at 20,000 x *g* for 15 min and resuspended in degassed buffer B (100 mM Na_2_PO_4_ [pH 7.4], 200 mM NaCl, and 1 mM EDTA). Fluc was then incubated in the presence of a four- and sixfold excess of pre-mixed AF555 donor and AF647 acceptor fluorophores, respectively, and placed on a rotator overnight at 4°C. Following the coupling reaction, excess dye was removed by at least two passes over separate gel filtration columns using a 7000 Da molecular weight cutoff Zeba Spin Desalting column (Thermo Fisher Scientific, USA) equilibrated in 50 mM tris (pH 7.5) supplemented with 20% (v/v) glycerol. The concentration and degree of labeling were calculated by ultraviolet absorbance.

### Refolding assays

The ability of denatured Fluc to spontaneously refold to a native state was assessed by measuring the return of enzymatic activity after denaturation. Denatured Fluc was prepared by incubation of native Fluc in unfolding buffer (50 mM Hepes-KOH [pH 7.5], 50 mM KCl, 5 mM MgCl_2_, 2 mM dithiothreitol [DTT], and 5 M GdHCl) for 30 min at room temperature. Spontaneous refolding of denatured Fluc was initiated by a 1:100 dilution into refolding buffer (50 mM Hepes-KOH [pH 7.5], 80 mM KCl, 5 mM MgCl_2_, 2 mM DTT, and 0.05% [v/v] Tween-20) such that the final concentration of Fluc was 10 nM. Refolding reactions with bacterial chaperones (i.e., KJE system) were left to incubate at 25°C for up to 120 min. Throughout the refolding reactions, aliquots were taken at various times and dispensed into the bottom of a white 96-well Costar plate (Sigma-Aldrich, USA). The luminescence reaction was initiated following injection of a 10-fold excess of assay buffer (25 mM glycylglycine [pH 7.4], 0.25 mM luciferin, 100 mM KCl, 15 mM MgCl_2_, and 2 mM ATP) into a single well, and 5 s after injection, the luminescence was measured for 10 s using a POLARstar Omega (BMG Labtech, Germany) plate reader. The injection and measurement procedure described above was then performed sequentially for each well to ensure consistency between the measurements. All measurements were performed at 25°C with the gain set to 4000. Refolding yields were calculated by normalizing to the activity of native (non-denatured) Fluc, which was treated as described above with the exception that GdHCl was omitted from the unfolding buffer. Chaperone-assisted refolding reactions were performed as described above with the exception that denatured Fluc was diluted into refolding buffer containing chaperones and nucleotide. Briefly, refolding assays containing chaperones were performed with Fluc in the presence of 0.6 μM DnaJ, 20 μM DnaK, 1.2 μM GrpE (when present) and 10 µM HtpG (when present) supplemented with 5 mM ATP unless otherwise specified.

### Total internal reflection fluorescence (TIRF) microscopy

#### Microscopy setup

Samples were imaged using a custom-built total internal reflection fluorescence (TIRF) microscopy system constructed using an inverted optical microscope (Nikon Eclipse TI-2) that was coupled to an electron-multiplied charge-coupled device (EMCDD) camera (Andor iXon Life 897, Oxford Instruments, UK). The camera was integrated to operate in an objective-type TIRF setup with diode-pumped solid-state lasers (200 mW Sapphire; Coherent, USA or Stradus 637–140, Vortran Laser Technology, USA) emitting circularly polarized laser radiation of 532 nm continuous wavelength. The laser excitation light was reflected by a dichroic mirror (ZT405/488/532/640; Semrock, USA) and directed through an oil-immersion objective lens (CFI Apochromate TIRF Series 60x objective lens, numerical aperture = 1.49) and onto the sample. Total internal reflection was achieved by directing the incident ray onto the sample at the critical angle (θ_c_) of ~67° for a glass/water interface. The evanescent light field generated selectively excites the surface-immobilized fluorophores, with the fluorescence emission passing through the same objective lens and filtered by the same dichroic mirror. The emission was then split using a T635lpxr-UF2 dichroic (Chroma, USA), passed through ET690/50m and ET595/50m (Chroma, USA) cleanup filters and the final fluorescent image projected onto the EMCDD camera. The camera was running in frame transfer mode at 5Hz, with an electron multiplication gain of 700, operating at −70°C with a pixel distance of 160 nm (in sample space).

#### Coverslip preparation and flow cell assembly

Coverslips were passivated as previously described ^23^, with minor modifications to functionalize the surface with biotin for protein immobilization. Briefly, 24 × 24 mm glass coverslips were cleaned by alternatively sonicating in 100% ethanol and 5 M KOH for a total of 2 h before aminosilanization in 2% (*v/v*) 3-aminopropyl trimethoxysilane (Alfa Aesar, USA) for 15 min. NHS-ester methoxy-polyethylene glycol, molecular weight 5 kDa (mPEG) and biotinylated-mPEG (bPEG; LaysanBio, USA), at a 20:1 (*w/w*) ratio, was dissolved in 50 mM 4-morpholinepropanesulfonic acid (pH = 7.4) buffer and sandwiched between two activated coverslips for a minimum of 4 h for initial passivation in a custom-made humidity chamber. PEGylated coverslips were then rinsed with milli-Q and PEGylated again as described above for overnight (~20 h) passivation. PEGylated coverslips were rinsed with milli-Q water, dried under nitrogen gas, and stored at −20°C until required. Prior to use, neutravidin (0.2 mg/ml; BioLabs, USA) in milli-Q was incubated on the passivated coverslip for 10 min to bind to the bPEG. Neutravidin functionalized coverslips were then rinsed with milli-Q, dried under nitrogen gas and adhered to a polydimethylsiloxane flow cell for use in single-molecule experiments. Finally, to reduce the non-specific binding of proteins to the coverslip surface, each channel in the microfluidic setup was incubated in the presence of 2% (v/v) Tween-20 for 30 min as previously described ^24^ and then washed with imaging buffer (refolding buffer supplemented with 6 mM 6-hydroxy-2,5,7,8-tetramethylchroman-2-carboxylic acid [Trolox]).

#### Surface immobilization of labeled proteins and acquisition of smFRET data

For smFRET experiments, labeled Fluc constructs were specifically immobilized to a neutravidin-functionalized and Tween-20–coated coverslip. To do so, labeled proteins (~ 50 pM final concentration) were diluted in imaging buffer and incubated in the flow cell for 5 min. These conditions would typically give rise to ~200 to 300 FRET-competent molecules per 100 μm^2^. Unbound proteins were then removed from the flow cell by flowing through imaging buffer.

All data were acquired using the TIRF microscope setup previously described following sample illumination using a 532-nm solid-state laser with excitation intensity of 2.6 W/cm^2^, and the fluorescence of donor and acceptor fluorophores was measured every 200 ms at multiple fields of view. An oxygen scavenging system consisting of 5 mM protocatechuic acid and 50 nM protocatechuate-3,4-dioxygenase was included in all buffers before and during image acquisition to minimize photobleaching and fluorophore blinking.

### Single-molecule fluorescence resonance energy transfer (smFRET) workflow

To benchmark the Fluc sensors for refolding experiments, we measured the FRET distributions of native and misfolded Fluc, the latter generated by removing 4 M guanidine hydrochloride from immobilized Fluc using imaging buffer. Next, Fluc was incubated with DnaJ (0.6 µM) and DnaK (20 µM) for 10 min to generate fully saturated Fluc-DnaK complexes that act as a starting point for each refolding experiment. Image acquisition was initiated, and the indicated molecular chaperones were flown into the microfluidic channel after 10 s (1 mL/min, 5 s dead-time) and the donor and acceptor fluorescence was measured at 10 fields of view for 6 min each (60 min total imaging time) and used to calculate the FRET efficiency. Unless otherwise specified, the concentrations of each molecular chaperone were as follows; DnaJ (0.6 µM), DnaK (20 µM), GrpE (1.2 µM), HtpG and variants (10 µM). For experiments in which the DnaK-inhibitor telaprevir (100 µM) or HtpG-inhibitor radicicol (60 µM) were present, the inhibitors were pre-incubated with the molecular chaperone combinations for 5 min prior to the addition of ATP to ensure effective binding. In each instance, the concentration of DMSO was < 5% and did not affect refolding kinetics. All experiments were performed in imaging buffer at room temperature.

### smFRET analysis

#### Molecule selection and FRET calculations

Single-molecule time trajectories were analyzed in MATLAB using the MASH-FRET user interface (version 1.2.2, accessible at https://rna-fretools.github.io/MASH-FRET/) ^25^. The approximate FRET value is measured as the ratio between the acceptor fluorescence intensity (*I*_Acceptor_) and the sum of both donor (*I*_Donor_) and acceptor fluorescence intensities after correcting for crosstalk between donor and acceptor channels. The formula for calculating the FRET efficiency is given by Equation 1, whereby the corrected acceptor intensity (denoted as *CI*_Acceptor_) is equal to *I*_Acceptor_*–* (g * *I*_Donor_) and g is the crosstalk correction constant. g is calculated as the ratio of fluorescence measured in the acceptor and donor detection channels following direct excitation of a protein labeled with a single donor fluorophore.

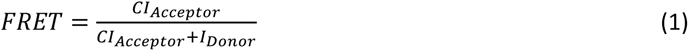

Briefly, donor and acceptor fluorescence channels were aligned following a local weighted mean transformation of images of TetraSpeck fluorescence beads and donor and acceptor fluorescence spots colocalized to identify FRET pairs. Molecules that displayed clear donor and/or acceptor photobleaching events or demonstrated anti-correlated changes in donor and acceptor fluorescence intensity were selected for subsequent analysis. The number of photobleaching events observed was used to determine the number of fluorophores present; only molecules in which a single donor and acceptor photobleaching event was observed were used for further analysis.

#### Trace processing and hidden Markov Model fitting

Selected molecules were denoised in MASH-FRET using the non-linear (NL) filter, which has been described previously ^26^, to accurately identify and quantify transitions between different FRET states during downstream processing. Parameter values were as follows; exponent factor for predictor weight, 5: running average window size, 1: factor for predictor average window sizes, 5. Data were truncated to only include FRET values acquired before donor or acceptor photobleaching. FRET efficiency data were exported to the state finding algorithm vbFRET (version vbFRET_nov12, https://sourceforge.net/projects/vbFRET/) and trajectories fit to a hidden Markov Model (HMM) to identify discrete FRET states, their residence times and the transition distributions between them. Default vbFRET settings were employed to fit data to the HMM, with the exception that the *mu* and *beta* hyperparameters were changed to 1.5 and 0.5, respectively, to prevent over-fitting.

#### Kinetic analysis of HMM fits

The HMM fits of individual FRET trajectories were further analyzed to extract key kinetic information arising from changes in Fluc conformation. To investigate transitions of interest, each transition (as determined from the HMM analysis) was sorted into different directional classes denoted generally as T_Before-After_, whereby ‘before’ refers to *F*_Before_ and ‘after’ refers to *F*_After_. For simplicity, FRET data were binned according to whether *F*_Before_ or *F*_After_ was > 0.3 (high) or < 0.3 (low), unless otherwise indicated, and thus, two different transition classes are possible: T_high-low_ and T_low-high_. The residence time (defined as the time that a molecule resides at *F*_Before_ prior to transition to *F*_After_) for each transition within a transition class was calculated and presented as cumulative histograms or the mean ± SEM. Since the FRET data could be well described as a two-state system in the context of chaperone binding (i.e., bound or non-bound), the cumulative residence times (determined as the period of time that the HMM fit resides at low-FRET states prior to a T_low-high_ transition, and vice-versa for T_high-low_ transitions) are presented. DnaK-release events are defined as a T_low-high_ transition, whereby the FRET state following a DnaK release event is equal to *F_after_* based on the HMM fits. The FRET state following DnaK release could then be shown as a violin plot or as a cumulative distribution. Since it is not possible to determine for how long a particular FRET state would have existed if not truncated due to photobleaching, the last measured FRET state was deleted and thus excluded from residence time calculations. Since data were smoothed during denoising, residence times shorter than that given by Equation 2 were not considered for further analysis. *F* is the imaging frame rate in milliseconds and *N_FA_* is the number of frames that were averaged during trace denoising.

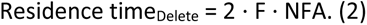

The residence times were fit with a double-exponential model and the two rates (*k_1_* and *k_2_*) and the contribution of each rate to the final fit determined. The accuracy of the fitting was validated using a bootstrapping approach, where residence times are sampled, fit to a double-exponential and the mean and standard error of *k_1_* and *k_2_* was determined by taking the mean distribution of fit values from each bootstrap attempt (n = 500). Finally, to determine whether there are changes in FRET immediately prior to or after chaperone dissociation, all T_low-high_ transitions were identified (which are indicative of chaperone dissociation), filtered such that only transitions with a residence time > 10 s were selected, and the FRET efficiency 10 s prior to and after each T_low-high_ was plotted. The FRET efficiency of all datapoints for each molecule was collated and are presented as FRET efficiency kernal density estimate (KDE) distributions or heatmaps.

### Modeling of smFRET data

To determine if experimentally observed smFRET data could be explained by a model in which DnaK dynamically transitions between different DnaK-bound states, theoretical simulations were performed. The observed DnaK-association rate was estimated by fitting the decrease in FRET efficiency post-DnaK-release (for Fluc^IDS1^ in the absence of HtpG, i.e., KJE; described above) to a single exponential, yielding a rate constant of *k_on_* = 0.55 s^−1^. Since it was hypothesized that *k_1_* from double-exponential fits of T_low-high_ residence times more accurately reflects true DnaK-dissociation times (see above), the average *k_1_* across all Fluc sensors was used to approximate the DnaK dissociation rate (*k_off_* = 0.15 s^−1^).

Using these *k_on_* and *k_off_* values, we simulated transitions between different DnaK-bound states (denoted as *state_n_*, where *n* is the number of bound DnaK proteins) every 1 s over a total of 360 s, consistent with the timescales of smFRET acquisitions. Based on the entropic-pulling model of Hsp70 function ^27,28^, we assumed that at least two or more DnaK proteins are required for full conformational expansion of the client. Accordingly, FRET states were assigned as follows: low-FRET when ≥2 DnaK proteins were bound, and high-FRET when one or none were bound. T_low-high_ residence times were then extracted from >500 simulated FRET trajectories as previously described for smFRET data, and the resulting distributions were plotted as cumulative density plots and fit using either single- or double-exponential models. To assess whether the number of potential DnaK-binding sites influences the simulated residence times, simulations were repeated with increasing maximum numbers of DnaK-bound states (*state_max_*). The average T_low-high_ residence time and the proportion of time spent in each *state_n_* were also determined as a function of *state_max_*.

### Statistical analysis

When comparing the means between multiple treatments, data was statistically analyzed using a one-way analysis of variance (ANOVA) with a Tukey’s multiple comparisons *post-hoc* test, with *P* ≤ 0.05 determined to be statistically significant. To assess whether the distribution of FRET states following DnaK release differed between treatments, a two-sample Kolmogorov-Smirnov test was performed for each pairwise comparison (with P < 0.05 considered statistically significant). All data analysis and presentation were performed using custom-written scripts on Python software or using GraphPad Prism 9 (GraphPad Software Inc; San Diego, USA). All original code for the analysis and visualization of smFRET data has been deposited at Zenodo at the following DOI: XXXXXXXX

### Code availability

All data needed to evaluate the conclusions in the paper are present in the paper and/or the Supplementary Materials. All unique reagents generated in this study are available from the lead contact without restriction. All original code has been deposited at Zenodo at the following DOI: https://doi.org/10.5281/zenodo.16791519, using version 1.0.3. Further information and requests for resources and reagents should be directed to and will be fulfilled by the lead contact, H.E.

## Results

### HtpG prevents DnaK-mediated inhibition by blocking client rebinding by DnaK and promoting controlled folding events

First, we established that the bacterial Hsp90 system (consisting of Dna<u>K</u>, DnaJ, Grp<u>E</u> and Htp<u>G</u>, denoted as KJEG) could refold a denatured client protein. For these experiments, we employed a previously reported firefly luciferase mutant (denoted as Fluc^IDS1^, since this protein can act as an interdomain sensor of protein folding via smFRET) ^11^. Fluc^IDS1^ refolding was monitored by measuring the luminescence from correctly folded and native protein over time. Fluc^IDS1^ was unable to return to an enzymatically active state except in the presence of the complete KJEG system (Fig 1A); notably, the inability of the KJE system to refold Fluc^IDS1^ is due to the high (yet physiologically relevant) concentrations of DnaK used in this assay ^11,12^. Crucially, Fluc refolding was dependent on GrpE-mediated handover of the client to HtpG since omission of either component prevented refolding.

**Fig 1:**
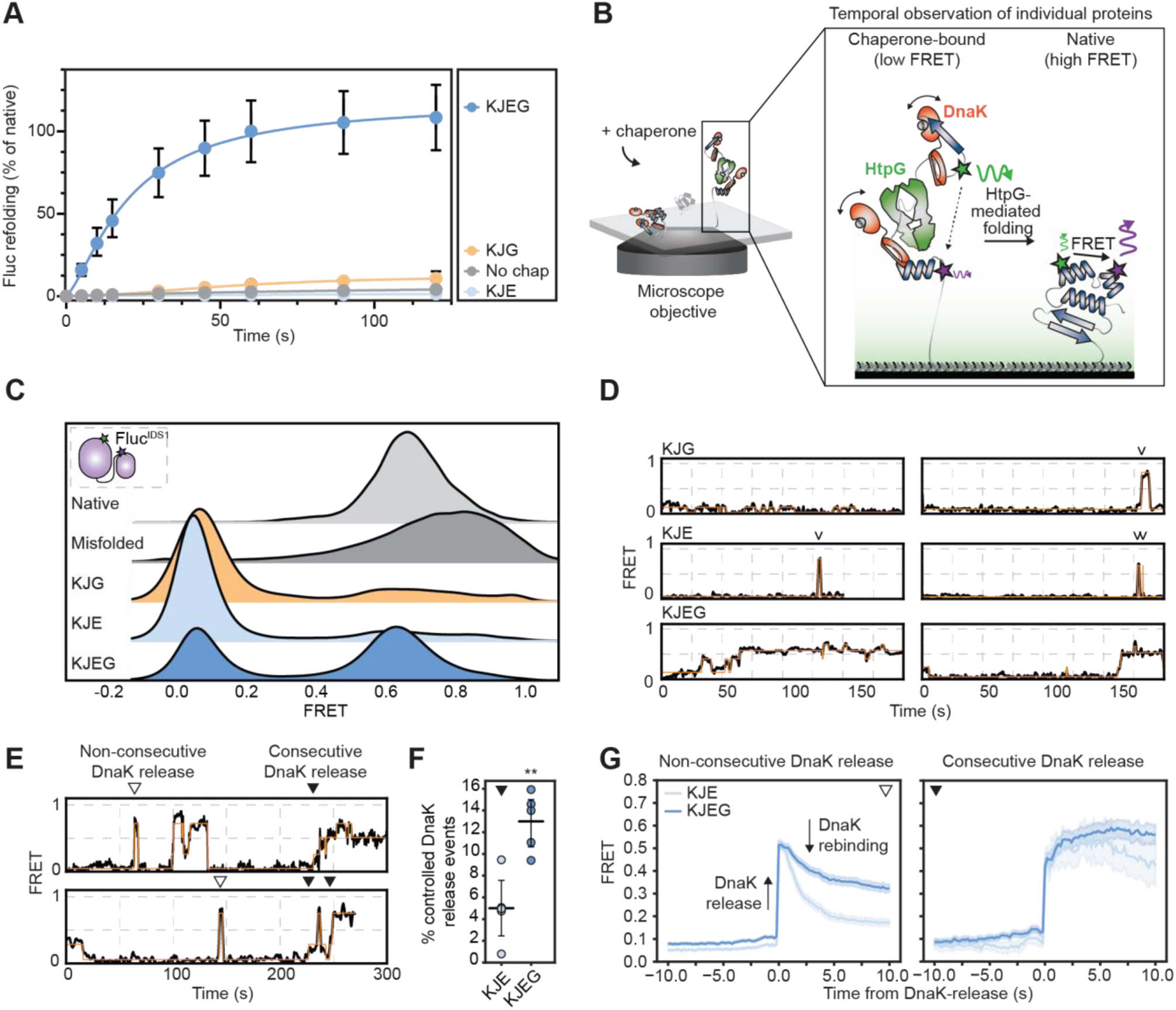
HtpG prevents the DnaK-mediated inhibition of client refolding by promoting controlled folding events. **(A)** Luciferase refolding assay in the absence or presence of the KJEG system. Fluc^IDS1^ (10 nM final concentration) was diluted 100-fold into refolding buffer alone (i.e., no chap) or supplemented with various combinations of molecular chaperones. Data shown represents the mean ± SEM from three independent experiments. **(B)** Schematic of the smFRET imaging setup. AF555/AF647-labeled Fluc^IDS1^ was immobilized to a neutravidin-functionalized coverslip and was illuminated by a 532-nm laser that selectively excites the AF555 donor fluorophore. The fluorescence from both AF555 and AF647 (acceptor) fluorophores was measured over time in the absence or presence of chaperones, and the FRET efficiency was calculated. **(C)** FRET efficiency kernel density estimate (KDE) distributions of Fluc^IDS1^ (*inset, relative position of FRET-pair is denoted*) in the native state, misfolded or during 60 min of incubation with the indicated combination of chaperones. Data is compiled from at least 193 individual trajectories for each treatment. **(D)** Example FRET trajectories from individual Fluc^IDS1^ molecules upon incubation with the indicated combination of molecular chaperones (HMM shown in orange). Arrows indicate the presence of ‘spike’ events. **(E)** Example FRET trajectories with DnaK-release events (i.e., transitions from < 0.3 to > 0.3 FRET) classified into either ‘non-controlled’ (*empty triangles*) or ‘controlled’ events (*filled triangles*) denoted. **(F)** The percentage of DnaK-release events classified as ‘controlled’. Data is presented as mean ± SEM. ** denotes p < 0.01 following an unpaired t-test. **(G)** The average FRET efficiency prior to and after ‘non-controlled’ (*left*) and ‘controlled’ (*right*) DnaK-release events ± SEM. Data is compiled from at least 1242 events per treatment.

Having established that the KJEG system can refold Fluc^IDS1^ to an enzymatically active state, we next investigated how the conformation of individual Fluc^IDS1^ proteins is altered during HtpG-mediated refolding using a combination of TIRF microscopy and smFRET (Fig 1B). Native Fluc^IDS1^ exhibited a single FRET distribution centred at ~ 0.6, which increased to ~ 0.8 upon formation of a chemically-induced misfolded state (Fig 1C), consistent with our previous work ^11^. Incubation of misfolded Fluc^IDS1^ with DnaJ/DnaK in the absence of either GrpE or HtpG, which was previously shown to inhibit refolding (Fig 1A), resulted in the formation of an ultra-low FRET state (centered at ~0.1). We have previously shown that this state is characteristic of DnaK-bound Fluc^IDS1^, whereby DnaK binding results in substantial conformational expansion of the client to resolve misfolded states ^10,11^. Analysis of individual Fluc^IDS1^ molecules from these treatments revealed that the DnaK-bound state was extremely long-lived but would occasionally exhibit rapid FRET ‘spikes’, indicative of DnaK dissociation (i.e., an increase in FRET) followed by rapid rebinding (i.e., a decrease in FRET) (Fig 1D, Fig S1A). These data are consistent with Fluc^IDS1^ becoming ‘trapped’ within the DnaK-bound state, which is not permissible for productive refolding. Refolding of Fluc^IDS1^ to a native-like FRET state was only observed in the presence of the complete KJEG system, with gradual transitions away from the ultra-low FRET (DnaK-bound) state towards a native-like FRET efficiency often occurring gradually, consistent with controlled folding of the client. Such controlled folding events, during which hydrophobic regions are potentially shielded to form the hydrophobic core of the protein, may explain how HtpG can assist Fluc^IDS1^ to escape rebinding by DnaK.

To interrogate this further, we classified DnaK-release events (i.e., transitions from < 0.3 to > 0.3 FRET) as being either those that exhibit consecutive increases in FRET (which we defined as ‘controlled’ or ‘gradual’) or those in which rebinding occurs after DnaK-release (defined as ‘non-consecutive’ or ‘non-controlled’, Fig 1E). Interestingly, there was a ~ 2.5-fold higher probability of controlled folding events when HtpG was present compared to when HtpG was omitted (Fig 1F). Plotting the average FRET efficiency prior to and after these DnaK-release events (Fig 1G) shows that, for non-consecutive events, the FRET efficiency decreased rapidly in the absence of HtpG (indicative of Fluc^IDS1^ misfolding and subsequent rebinding by DnaK); however, the rate and amount of DnaK rebinding was substantially less in the presence of HtpG (Fig 1G). This could be due to either HtpG-mediated refolding being more likely to result in the client adopting a native-like fold, or HtpG preventing DnaK rebinding, or a mixture of both. Fitting the DnaK-rebinding curves to a single exponential supports it being a mixture of both mechanisms (Fig S1B-C) since HtpG reduces both the rate constant (indicating slower DnaK rebinding rates) and the total amplitude (indicating reduced DnaK engagement due to native folding or HtpG-mediated blocking). Conversely, a gradual increase in FRET efficiency was observed following controlled DnaK-release events for all conditions tested (Fig 1G). Crucially, minimal DnaK rebinding was observed, indicating that controlled folding events, which are promoted by HtpG (Fig 1F), are more likely to result in the client reaching a native fold rather than non-controlled events.

### Controlled folding is driven by HtpG ATPase activity and coordination with DnaK

To interrogate how Hsp90 cooperates with the Hsp70 system to promote efficient client refolding, we performed smFRET experiments in the presence of a HtpG inhibitor (radicicol, which prevents ATP binding) or with HtpG mutants that have been previously reported to reduce ATP hydrolysis (i.e., HtpG^E34A^) or prevent interaction with DnaK (HtpG^R355L^) ^8^. As expected, the HtpG mutants or radicicol-inhibited HtpG could not refold Fluc^IDS1^ (Fig 2A, Fig S2A, example traces in Fig S2B) and had a reduced capacity to promote controlled folding events compared to wild-type HtpG (Fig 2B). Interestingly, HtpG^E34A^ promoted higher rates of controlled folding events compared to radicicol-treated HtpG, indicating that ATP binding alone (or even slow rates of ATP hydrolysis) is important in facilitating controlled folding events (Fig 2B). As before, these controlled DnaK-release events resulted in minimal DnaK rebinding, again suggesting that these events promote correct folding of the client (Fig 2C). This indicates that only a functioning HtpG system can reduce DnaK re-engagement following non-controlled DnaK release events. We next investigated whether HtpG regulates DnaK-release events to promote optimal client refolding by looking at the first FRET state of an individual client molecule immediately following DnaK-release (Fig 2D, based on HMM fits). A distribution analysis revealed that HtpG significantly reduces transitions to typically misfolded high-FRET Fluc^IDS1^ structures (i.e., > 0.6) compared to when ATP binding/hydrolysis is inhibited (i.e., with HtpG^E34A^ or radicicol, Fig 2E). This finding is consistent with previous experiments that suggest ATP hydrolysis is critical in ‘controlling’ client refolding ^29^. Interestingly, transitions to high-FRET states following DnaK-release were more common in the presence of HtpG^R355L^ compared to other treatments, even those that did not contain HtpG (Fig 2E), suggesting that indirect interactions between DnaK and HtpG reduce collapse of Fluc^IDS1^ into misfolded structures. Wild-type HtpG also accelerates the rate of observed DnaK-release events (observed as a decrease in the time spent in a DnaK-bound state) compared to the HtpG mutant isoforms tested (Fig S2C), whereby it increases both the occurrence and folding efficiency of Fluc^IDS1^ folding events. Collectively, these data suggest that, through direct interactions with DnaK, HtpG utilizes ATP hydrolysis to increase the probability of Fluc^IDS1^ folding correctly by actively protecting it from forming non-productive misfolded intermediates following release by DnaK.

**Fig 2:**
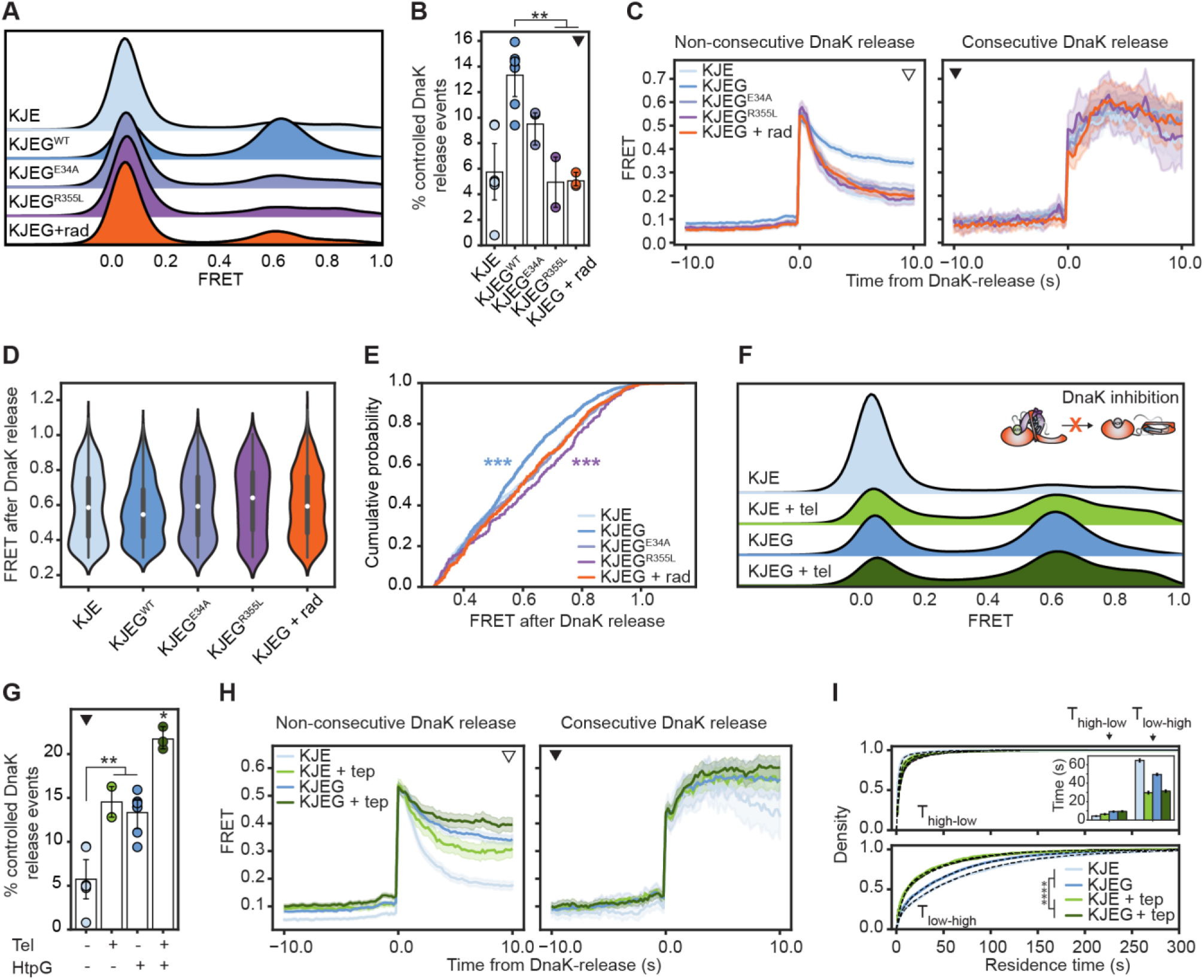
HtpG selectively prevents DnaK rebinding during folding events. **(A)** FRET efficiency KDE distributions of Fluc^IDS1^ during 60 min of incubation with the indicated combination of chaperones/inhibitors (60 µM radicicol). Data is compiled from at least 686 individual trajectories for each treatment. **(B)** The percentage of DnaK-release events that are classified as ‘controlled’. **(C)** The average FRET efficiency prior to and after ‘non-controlled’ (*left*) and ‘controlled’ (*right*) DnaK-release events. Data is compiled from at least 431 events per treatment. **(D)** The distribution of Fluc^IDS1^ FRET states immediately following DnaK-release used for distribution analysis in panel E. **(E)** Cumulative histogram of the FRET state immediately following DnaK-release. Data were analyzed using a Kolmogorov–Smirnov test for differences in the distribution. *** denotes statistically significant difference (p< 0.001) between the indicated treatment and the KJE, KJEG^E34A^ and KJEG + radicicol conditions. **(F)** FRET efficiency KDE distribution of Fluc^IDS1^ during 60 min of incubation with the indicated combination of chaperones/inhibitors (100 µM telaprevir). Data is compiled from at least 1207 individual trajectories for each treatment. **(G)** The percentage of DnaK-release events that are classified as ‘controlled’. **(H)** The average FRET efficiency prior to and after ‘non-controlled’ (*left*) and ‘controlled’ (*right*) DnaK-release events ± SEM. Data is compiled from at least 432 events per treatment. **(I)** Cumulative residence time data showing the time Fluc^IDS1^ remains in a DnaK-bound state (i.e., < 0.3 FRET) prior to a non-bound state (i.e., T_low-high_) and *vice-versa*. The average residence time (*inset*) is calculated and presented as mean ± SEM for clarity. For panels B, G and I, a one-way ANOVA with Tukey’s post-hoc test was performed, with *, ** and *** denoting statistical significance of p < 0.05, 0.01 and 0.001, respectively.

### HtpG reduces DnaK rebinding to folding intermediates but not misfolded clients

Since our data suggests that HtpG prevents DnaK-rebinding during folding, we investigated whether optimal refolding could be recapitulated by reducing the binding activity of DnaK. Indeed, addition of the DnaK-specific inhibitor telaprevir to typically inhibitory concentrations of DnaK (Fig 1A) allowed Fluc refolding to be partially restored in the absence of HtpG and completely restored it in the presence of HtpG (Fig S3A). This suggests that (i) reducing DnaK-association improves folding and (ii) DnaK is not completely inhibited at these concentrations of telaprevir, consistent with previous work ^30^. Repeating these experiments at the single-molecule level yielded similar results, whereby the addition of telaprevir improved refolding yields in the absence of HtpG (Fig 2F, example traces in Fig S3B), but not during HtpG-mediated folding. Importantly, telaprevir resulted in a substantial increase in misfolded Fluc^IDS1^ for both conditions (Fig S3C). Interestingly, telaprevir-induced DnaK inhibition resulted in an increased propensity of controlled folding events relative to the non-inhibited controls (i.e., with or without HtpG, Fig 2G) and, as expected, reduced the rate of DnaK-rebinding after DnaK release (Fig 2H, Fig S3D-E). However, there are some notable observations; (i) HtpG is more effective than telaprevir at slowing DnaK-rebinding following its release (Fig S3E) and (ii) HtpG-induced inhibition of DnaK rebinding is primarily selective for Fluc^IDS1^ molecules following DnaK-release (since DnaK binding to misfolded Fluc^IDS1^ is not affected, as evidenced by a lower abundance of misfolded species; Fig S3C) whereas telaprevir broadly inhibits DnaK binding across all Fluc^IDS1^ states. This observation suggests that HtpG actively prevents DnaK rebinding while bound to Fluc^IDS1^, thus facilitating high DnaK association rates to misfolded clients without compromising the ability of the client to fold following DnaK release. Indeed, the addition of telaprevir also resulted in substantially faster rates of observed DnaK-release events compared to HtpG-mediated folding alone (Fig 2I). Since DnaK dissociation is independent of the rate of association ^12^, these data are consistent with previous reports that longer residence times (e.g., in the case of HtpG without telaprevir) correlate with an increased number of DnaK proteins bound to each client ^11^. As such, Fluc^IDS1^ is protected from misfolding in the presence of HtpG but still amenable to controlled folding processes. Together, these data support a mechanism whereby slower DnaK association rates, mediated either by HtpG or telaprevir, result in more ‘controlled’ folding events and improved folding outcomes, potentially since multiple asynchronous DnaK dissociation events can occur during domain folding without interference from DnaK rebinding.

### HtpG mediates controlled folding across multiple Fluc regions and is essential for establishing native interdomain contacts

Since Fluc^IDS1^ reports only on the distance between the N- and C-terminal domains, we developed additional cysteine mutants that allow us to observe the folding of other Fluc regions. To this end, FRET-pairs were positioned to generate another interdomain sensor that also reports on the domain linker (Fluc^K329C-K534C^; denoted as Fluc^IDS2^), an N-terminal domain sensor (Fluc^A12C-K329C^; denoted as Fluc^NDS1^) and a C-terminal domain sensor (Fluc^K378C-K496C^; denoted as Fluc^CDS1^) (Fig 3A). Refolding of all Fluc constructs was performed in the absence or presence of HtpG. Both the FRET efficiency and proportion of time that each molecule spends in an ultra-low (DnaK-bound) FRET state were determined and act as an inverse measure of the amount of native protein and thus refolding. In the absence of HtpG, both interdomain sensors (i.e., Fluc^IDS1^ and Fluc^IDS2^) remained predominantly DnaK-bound over time (Fig 3B-C), while the intradomain sensors could partially (in the case of Fluc^NDS1^) or completely (for Fluc^CDS1^) escape the DnaK-trapped state and return to more native-like FRET states (Fig 3B). However, in the presence of HtpG, the rate of refolding (determined by fitting the decay curve of DnaK-bound molecules, Fig 3C) was ~ 11-fold faster for Fluc^IDS1^, ~ 9-fold for Fluc^IDS2^, ~ 4.5-fold for Fluc^NDS1^ and ~ 2-fold for Fluc^CDS1^ (Fig 3D), with all sensors returning to native-like FRET efficiencies. Interestingly, HtpG did not alter the order in which each Fluc domain folded; from fastest to slowest this was Fluc^CDS1^ > Fluc^NDS1^ > Fluc^IDS2^ > Fluc^IDS1^. Indeed, after 15 min, the interdomain regions probed by Fluc^IDS1^ and Fluc^IDS2^ were DnaK-bound (i.e., completely unfolded) ~ 40–60% of the time compared to only 15% for the N- and C-terminal domain sensors. While it has previously been suggested that the Hsp70-Hsp90 cascade increases refolding yields but not folding kinetics ^12^, the much faster rates observed here in the presence of HtpG likely reflect the relatively poor refolding ability of AviTagged Fluc, which necessitates a higher dependence on chaperone function for folding ^11^. Regardless, these data suggest that (i) interdomain folding comes with more kinetic traps (and thus more critically necessitates the action of HtpG), (ii) the N- and C-terminal domains fold independently to prevent misfolding, (iii) the formation of native interdomain contacts is the final stage of folding and requires both N- and C-terminal domains to be mostly (if not completely) folded, and (iv) HtpG accelerates the rate of refolding but does not change the order in which each domain folds.

**Fig 3:**
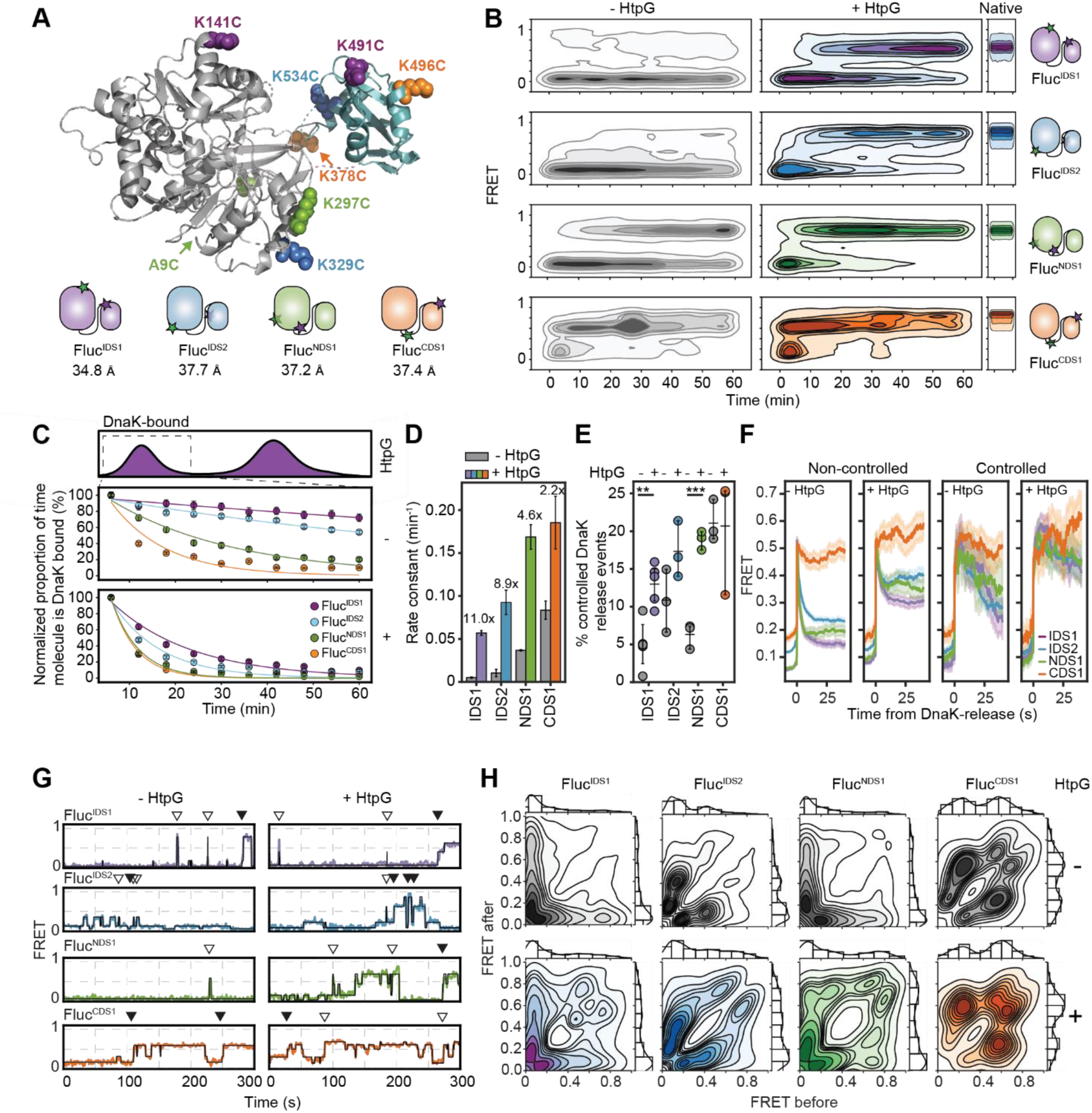
HtpG accelerates the folding of all Fluc domains and is critical for establishing native interdomain contacts. **(A)** Crystal structure of Fluc (ribbon; N-terminal domain [*gray*] and C-terminal domain [*blue*] are denoted, PDB:1LCI ^31^) with the position of introduced cysteines for each smFRET sensor denoted (*spheres*). Cartoon representation of the relative position and distance (in Å) of each FRET-pair on Fluc is shown below. **(B)** FRET intensity heatmaps for each Fluc sensor when incubated in the presence of the KJE system alone (-HtpG) or in the presence of HtpG (+ HtpG). Native Fluc is shown on the right. Data is compiled from at least 1252 individual trajectories for each treatment. **(C)** The proportion of time that each Fluc protein is DnaK-bound (i.e., < 0.3 FRET). Calculated from data presented in panel B. Data for each treatment was fit with an exponential decay curve. **(D)** The rate of refolding of each Fluc sensor derived from the rate constants calculated from the exponential fit in C. The fold increase in refolding rate when HtpG is present is denoted. **(E)** The percentage of DnaK-release events that are classified as ‘controlled’ when refolded by the KJE system alone or supplemented with HtpG. Each Fluc sensor in the absence or presence of HtpG was statistically analyzed by an unpaired t-test, with ** and *** denoting statistical significance of p < 0.05 and 0.01, respectively. Data comes from at least two biological repeats. **(F)** The average FRET efficiency prior to and after ‘non-controlled’ (*left*) and ‘controlled’ (*right*) DnaK-release events ± SEM. Data is compiled from at least 115 events per treatment. **(G)** FRET trajectories for each Fluc sensor when incubated in the presence of the KJE system only or when supplemented with HtpG. The presence of ‘non-controlled’ (*open triangle*) or ‘controlled’ (*closed triangle*) DnaK-release events are denoted above the traces. **(H)** Transition density plots (TDPs) of each Fluc sensor when incubated in the presence of the KJE system only (*top*) or when supplemented with HtpG (*bottom*).

As was observed for Fluc^IDS1^, HtpG increased the proportion of ‘controlled’ DnaK-release events within the N-terminal domain (i.e., Fluc^NDS1^) and trended higher for Fluc^IDS2^ (Fig 3E); notably, there was no effect within the C-terminal domain (i.e., Fluc^CDS1^), which is the region of Fluc that is least dependent on chaperone intervention for refolding ^10^. Consistent with observations from Fluc^IDS1^, DnaK-rebinding following non-controlled DnaK-release (as indicated by the decrease in FRET efficiency) was substantially less for all Fluc sensors when in the presence of HtpG (Fig 3F), except for Fluc^CDS1^. Interestingly, the FRET efficiency prior to non-controlled DnaK release events (i.e., those most likely to result in misfolding and DnaK-rebinding) was lower for Fluc^IDS1^ and Fluc^NDS1^ (~ 0.05) in the absence of HtpG compared to when HtpG was present (~ 0.08, Fig 3F, Fig S4A), which suggests that refolding from completely expanded states is non-productive for these regions of Fluc. While this difference in FRET efficiency is relatively small, due to the non-linear (inverse sixth-power) relationship between donor-acceptor distance on FRET efficiency, a small change in FRET at these scales constitutes a substantial (~ 1 nm) difference in distance (Fig S4B). Notably, Fluc regions characterized by low FRET efficiencies prior to DnaK-release (Fig S4A) were correlated with longer T_low-high_ residence times (Fig S4C) and a higher dependence on HtpG to facilitate controlled folding (Fig 3E). Thus, these data indicate that HtpG reduces DnaK-mediated client expansion to prime these regions of Fluc for optimal folding. Moreover, controlled’ DnaK-release events resulted in less DnaK-rebinding for all Fluc sensors, especially in the presence of HtpG where minimal rebinding was observed even 40 s after DnaK-release (Fig 3F). Collectively, these data are consistent with HtpG promoting controlled folding events that result in more native-like conformations.

By filtering the data for those molecules that are DnaK-bound (i.e., visit FRET states < 0.3, which is a prerequisite for HtpG function ^32^), we interrogated how HtpG regulates transitions between different Fluc conformational states during productive folding. Individual FRET traces were fit with an HMM (Fig 3G, additional traces in Fig S4D) and the distribution of FRET transitions are represented as a function of the FRET state before and after each transition as a transition density plot (TDP, Fig 3H). In the absence of HtpG, Fluc molecules were often characterized by short-lived ‘spikes’ from an ultra-low-FRET DnaK-bound state to either intermediate states (~ 0.4 for Fluc^IDS2^) or high-FRET misfolded states (for Fluc^IDS1^ and Fluc^NDS1^); this is observed at the population level by high transition density along the TDP axes. Interestingly, Fluc^IDS2^ appeared to report on exchanges between two different DnaK-bound states (with FRET efficiencies of ~ 0.1 and 0.4, respectively). Since such low FRET states are not present in the absence of DnaK, it is likely that these may be due to local folding of the linker and/or C-terminal domain. This is supported by residence time data, whereby transitions away from the ultra-low-FRET DnaK-bound state occur at similar timeframes for Fluc^CDS1^ and Fluc^IDS2^ (Fig S4C). As such, all Fluc sensors are constrained within DnaK-bound states in the absence of HtpG (except for Fluc^CDS1^), whereby DnaK dissociation is more often followed by DnaK-rebinding than productive folding events.

Crucially, HtpG alleviated the DnaK-induced block for all Fluc sensors by promoting transitions from intermediate FRET states (~0.2–0.5) to more native-like conformations (~ 0.6–0.8), especially for Fluc^IDS2^ and Fluc^NDS1^ (Fig 3G), consistent with a higher proportion of controlled DnaK-release events (Fig 3F). At the population level, these are evident as increased transition density along the top axis of the TDPs (Fig 3H). Interestingly, Fluc^IDS2^ often dynamically transitioned between an intermediate FRET state (~0.6) and the native state (~0.8) (Fig S4D), with transitions to the native state increasing over time (Fig S4E). Additionally, transitions away from the native state were poorly tolerated at early timepoints (i.e., the FRET efficiency remained low and did not return to native-like states) while those at later stages of folding recovered faster (i.e., quickly returned to the native state, Fig S4F). Collectively, the transition of Fluc^IDS2^ between two distinct states (likely due to DnaK association/dissociation at a single binding site) suggest that the folding of other protein regions (e.g., Fluc^NDS1^ or Fluc^CDS1^) is a pre-requisite for stable Fluc^IDS2^ folding. Also, while there is a preference for stepwise domain folding (i.e., CTD > NTD > interdomain), DnaK dissociation is an intrinsically stochastic process. As such, it is likely that the conformational fluctuations observed within each Fluc sensor occur simultaneously within individual proteins and that folding of different domains/regions occurs while other regions are spatially separated to prevent aberrant misfolding events.

### Heterogenous DnaK binding drives region-specific conformational expansion and folding kinetics

When we further interrogated the distribution of DnaK-dissociation events (i.e., T_low-high_, Fig S4C), we found that the data were best described by a double-exponential with two rate constants (Fig S5A-B); generally, this featured a relatively conserved fast-rate (*k_1_*, ~ 0.1 – 0.25 s^−1^) and a slow rate (*k_2_*, ~ 0.01 – 0.03 s^−1^) (Fig S5C) for each Fluc sensor. For domain sensors with longer average residence times (i.e., Fluc^IDS1^ and Fluc^NDS1^), the slow rate was ~ 2-3-fold slower compared to those sensors with shorter residence times (i.e., Fluc^CDS1^ and Fluc^IDS2^) and accounted for a larger portion of the overall fit (>70% vs. 20–50%, respectively) (Fig S5C), indicating that this slow rate dominates the occurrence of Fluc refolding events for these sensors. Since DnaK dissociation is typically defined by a single exponential rate ^33,34^, we next explored why this “simple” on/off system was best described by a double exponential rate, and why different regions of Fluc exhibited different residence times under the same conditions.

Based on our smFRET data, we hypothesized that the low-FRET (< 0.3) state reflects a dynamic and heterogeneous ensemble of multiple DnaK molecules bound to a single Fluc client, consistent with the entropic pulling model of Hsp70 function in which simultaneous binding of several Hsp70 proteins help resolve misfolded states ^11,27^ (Fig 4A). Escape from this expanded low-FRET conformation likely occurs upon dissociation of at least one DnaK, leaving one or no DnaK subunits bound, which results in a more compact conformation (FRET > 0.3). To explore this, we performed kinetic simulations of DnaK binding to a single Fluc molecule, using a fixed association (*k_on_*) and dissociation rate (*k_off_*, as determined from smFRET experiments) to model the dynamic exchange between different DnaK-bound states. Based on predicted DnaK binding sites on Fluc ^35^ and the position of fluorophores for each sensor, we assumed a maximum of 2–5 potential binding sites per Fluc region and tracked the number of bound DnaK molecules over time. As expected, Fluc dynamically transitioned between different DnaK-bound states (Fig 4B), with Fluc spending a greater proportion of time bound by more DnaK molecules as the number of potential binding sites increased (Fig S5D). To see if this model reflects our experimental data, we assigned FRET values based on DnaK occupancy: low-FRET for two or more bound DnaK proteins, and high-FRET when one or none were bound (Fig 4C). This approach allowed us to test whether differences in residence times observed experimentally could arise from a heterogeneous ensemble of multiple bound states. Indeed, when only three DnaK-bound states were possible, the simulated residence times were short (~ 6 s) and fit well to a single exponential; conversely, increasing the number of possible DnaK-bound states resulted in an increased average residence time with a distribution best described by a double exponential (consistent with our experimental smFRET data) (Fig 4D). The simulated FRET trajectories also appeared similar to the experimental smFRET traces (Fig 4C) and produced bi-exponential rate constants comparable to experimentally determined rates for our Fluc sensors (*k_1_*, ~ 0.1 – 0.33 s^−1^; *k_2_*, ~ 0.008 – 0.046 s^−1^, Fig S5E). Interestingly, when the observed rate of DnaK-rebinding post-DnaK release for Fluc^IDS1^ was used to define *k_on_* in the simulations, only minor changes in residence times were observed (Fig S5F), consistent with experimental results (Fig S4C). Instead, the number of states was the most substantial determinant of residence times. As such, these data suggest that the number of potential DnaK-binding sites plays a critical role in regulating Fluc refolding and explains how different Fluc regions exhibit different residence times under the same experimental conditions. This potentially provides a mechanism by which the generic function of Hsp70 and/or Hsp90 (i.e., client binding and release) can be tailored to the specific needs of the client.

**Fig 4:**
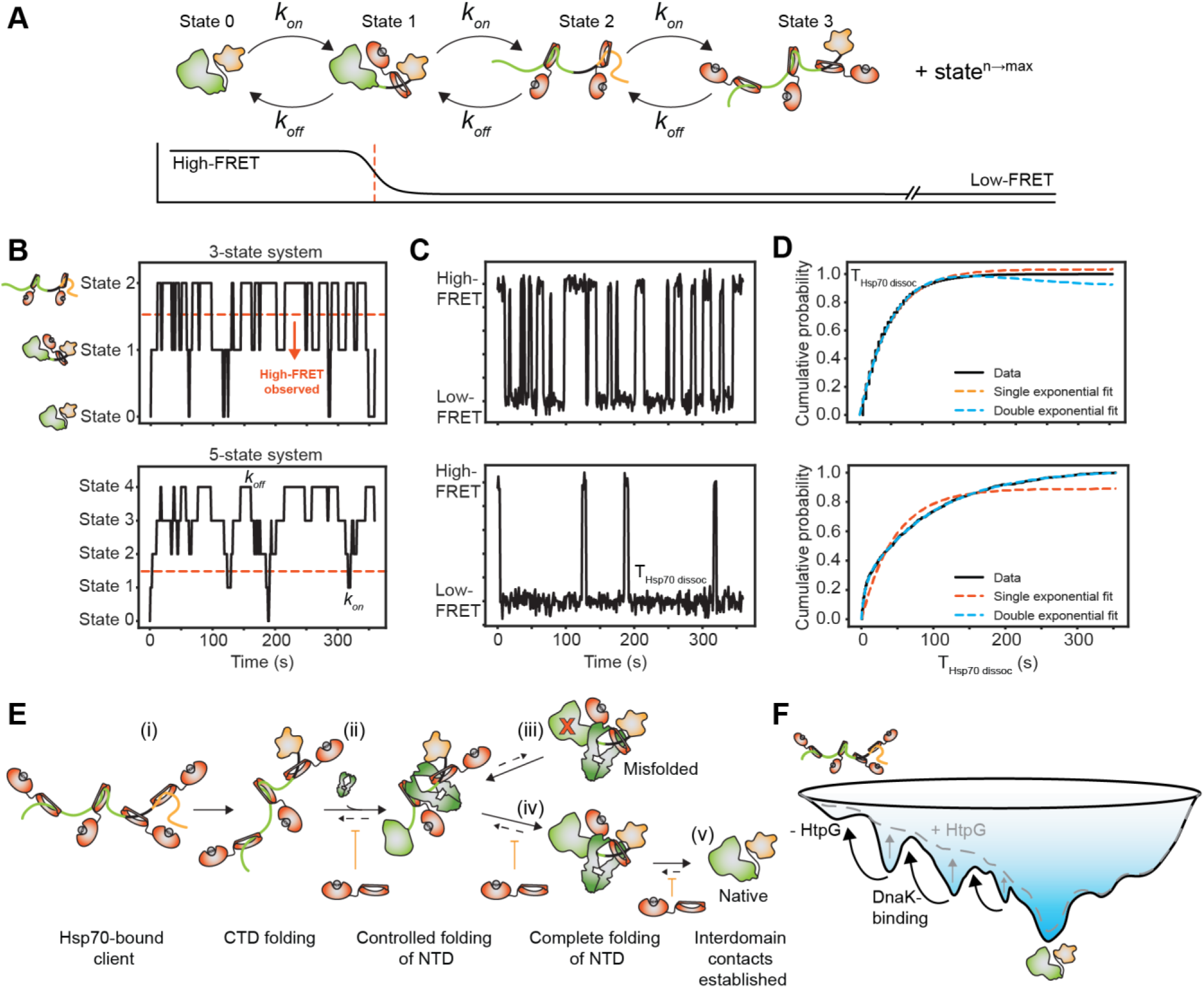
Dynamic exchange of DnaK between heterogeneous bound states regulates the kinetics of Fluc conformational expansion and folding. **(A)** Schematic illustrating the theoretical relationship between the number of client-bound DnaK molecules and the observed FRET efficiency of Fluc sensors. Since entropic pulling stipulates that binding of multiple Hsp70 subunits are required for full conformational expansion (i.e., low-FRET) ^28^, a threshold of < 2 bound-DnaK proteins to observe high-FRET states was defined (*orange line*). Transitions between different DnaK-bound states are defined by a single experimentally derived *k_on_* and *k_off_*. **(B)** Example simulated data, showing transitions between either three (*top*) or five (*bottom*) different possible DnaK-bound states. **(C)** FRET trajectories calculated from simulations in panel B, where transitions to the ‘high-FRET’ state only occur upon transitions to state 1 or lower. **(D)** The distribution of residence times from simulated FRET trajectories (panel C) were determined and plotted as cumulative density plots. Fits of the data to single- or double-exponential distributions are shown. **(E)** Model of HtpG-mediated folding of Fluc. (i) DnaK-mediated conformational expansion of Fluc occurs and is followed by (ii) folding of the C-terminal domain in an HtpG-independent manner. The folding of the N-terminal domain occurs next in an HtpG-mediated process, whereby (iii) HtpG utilizes ATP hydrolysis to promote controlled folding events, prevent DnaK rebinding and minimize misfolding, which is assisted by spatial separation of N- and C-terminal domains through the HtpG lumen. Should misfolding occur at any stage (*red X*), (iv) DnaK rebinding can still occur to resolve the misfolded state. (v) Upon independent folding of the N- and C-terminal domain, interdomain contacts are established in the final native state. **(F)** Mechanism of HtpG/DnaK-mediated folding. DnaK prevents the accumulation of misfolded client proteins in kinetic traps by returning them to a high entropy (i.e., conformationally expanded) conformation while HtpG smooths the folding landscape to the native state.

## Discussion

We set out to describe the conformation of an individual client protein as it is engaged by the bacterial Hsp70/Hsp90 chaperone systems during productive folding using smFRET. In doing so, we found that HtpG (i) reduces DnaK-rebinding during folding events, thereby allowing the client to escape from DnaK-induced inhibition, (ii) reduces client misfolding following DnaK-release and (iii) promotes ‘controlled’ folding of the client to the native state (Fig 4E). Collectively, HtpG acts to release the Hsp70-induced folding block when levels of DnaK prevent productive folding and smooth the folding landscape to increase the rate of client folding to the native state (Fig 4F).

Interestingly, HtpG increases productive folding of clients through enhancing thermodynamic stability and protecting against kinetic traps (as demonstrated by increased ‘controlled’ folding events and reduced misfolding following DnaK release), with only subtle contributions from changes in chaperone-induced conformational kinetics (e.g., DnaK-release and binding times). The improved thermodynamic stability is likely caused by a combination of factors. Firstly, HtpG/DnaK-mediated chaperone folding prevents interdomain misfolding by spatially separating the N- and C-terminal domains, which ensures that each domain can fold productively. This is consistent with previous reports that each Fluc domain can independently refold from expanded states ^36^, while misfolding occurs due to erroneous interdomain contacts. Indeed, interdomain misfolding is a general phenomenon observed for many multi-domain proteins and is often rate limiting for folding ^37,38^. Our smFRET and simulation data suggest that the number and position of Hsp70 binding sites facilitates efficient separation of domains to prevent misfolding (via additional sites at interdomain regions) while still enabling intra-domain folding events (via reduced binding sites). This effect is likely amplified upon Hsp90 binding, where client proteins are often threaded through its lumen ^20,39,40^, allowing each domain to fold independently on either side of the chaperone. Furthermore, domain refolding is more productive under conditions where external force is applied (as is the case when the client is DnaK-bound) since interdomain misfolding is reduced ^41^. As such, the client can fold down a smooth folding landscape to the native state without falling into kinetic traps (Fig 4F). Such a result can also explain why Hsp70 is involved during the early stages of protein folding (i.e., co-translationally on the ribosome), whereby interdomain misfolding is unlikely since the complete polypeptide has not emerged from the ribosome tunnel ^36,42^. Indeed, fewer Hsp70s are likely needed to resolve misfolded states during co-translational folding since the emerging polypeptide is tethered to the bulky ribosome, thereby enhancing the entropic pulling forces exerted by Hsp70 upon binding; such mechanisms have also been described in the context of protein translocation ^27^. Hsp90 therefore becomes more essential once the entire protein has been translated, whereby the number of available Hsp70 binding sites is much higher and Hsp90 is required to provide a surface whereby folding can occur productively without competition from Hsp70. Additionally, both smFRET and simulation data suggest that Hsp90-mediated folding is energetically less expensive compared to Hsp70-mediated folding, consistent with previous observations ^43^. Indeed, we suggest that the ‘holdase’ function of Hsp70 is somewhat of a misnomer (as it implies stable binding) and is energetically expensive, since continual and dynamic Hsp70 exchange is required to maintain the expanded bound state.

Secondly, HtpG promotes escape from DnaK-bound states by facilitating a 2–3-fold increase (for some domains) in controlled folding events, which are characterized by gradual or incremental FRET increases to stable, near-native states. These incremental FRET changes reflect progressive compaction events as Fluc adopts multiple folding intermediates following DnaK-release, which appear to be regulated by a combination of (i) ATP binding and hydrolysis by HtpG and (ii) the association/dissociation kinetics of DnaK. The mechanism by which Hsp90 utilizes ATP hydrolysis to promote client folding has been a longstanding and continuing debate in the field ^40,44,45^, however our data (and others) now suggest that it is critical in remodeling the client for folding ^21,29^. This is likely mediated by the substantial conformational rearrangements that HtpG undergoes upon ATP hydrolysis ^16^, whereby twisting of each monomer can remodel client binding sites in the Hsp90 lumen ^46,47^ and/or produce a substantial force on the client such that it can ‘slide’ through the lumen ^21^. Movement of the client through the lumen, albeit slight, potentially remodels the folding landscape of the protein by promoting local conformational compaction events that prevent misfolding. Indeed, optical-tweezer experiments have demonstrated that HtpG induces local chain compaction from an expanded client state following force release (which is mimicked in our system by DnaK-release) in an ATP-hydrolysis-dependent manner ^29^, supporting our conclusion that HtpG mediates ‘controlled’ folding upon ATP hydrolysis. Recent cryo-EM data with eukaryotic Hsp90 suggest that both client binding and ATP hydrolysis promote Hsp70 dissociation ^20,48^, providing a mechanism by which Hsp90 synchronizes its force-driven folding activity with DnaK release, allowing the client to fold in a controlled manner with access to previously shielded regions. This is likely assisted by the asymmetric nature of ATP hydrolysis on HtpG (i.e., hydrolysis can occur on either subunit) ^20,49,50^, whereby stepwise remodeling and release of the client promotes folding. Lastly, HtpG has been suggested to provide a more hydrophilic surface (compared to DnaK) for folding to occur, which might contribute to the stepwise or gradual compaction of the client observed in this work ^51^.

We were also able to show that controlled folding of the client is highly dependent on DnaK association and dissociation rates. This is largely consistent with a suite of ensemble-based assays, whereby high-concentrations of Hsp70 are inhibitory but can be recovered by either reducing the concentration of Hsp70 ^12^, inhibiting Hsp70 (e.g., with telaprevir, this work) or increasing NEF-mediated Hsp70 dissociation ^11^. The reduced rate of DnaK-rebinding (mediated by HtpG or telaprevir) increases opportunities for NEF-mediated DnaK dissociation events to be kinetically synchronized, thus allowing the formation of intradomain contacts that would otherwise not occur if bound DnaK was present. This also has the effect of producing smaller conformational rearrangements from the expanded state, facilitating the formation of a folding ‘nucleus’ from which additional folding events can occur productively with reduced risk of misfolding. Whilst the folding of individual clients via asynchronous DnaK release has been proposed previously ^10,52–54^, here we have been able to temporally observe it for single client molecules.

Our experimental data is also consistent with kinetic simulations, whereby decreasing DnaK association rates and/or increasing DnaK dissociation rates shifts the binding equilibrium to lower-bound states, thereby increasing opportunities for multiple dissociation events/compaction events to occur. In this instance, optimal folding likely occurs at the transition point between DnaK-mediated conformational expansion (to resolve misfolded states) and compaction of the client (to promote folding). HtpG enhances this process by reducing DnaK association to the folding client, either by promoting more native-like states (with less exposed hydrophobicity) via ATP-hydrolysis-mediated controlled folding, or by sterically occluding DnaK from rebinding. It is also possible that DnaK-mediated folding (via client association/dissociation) can occur while bound to ATP-bound HtpG, especially since ATP hydrolysis is rate limiting for the HtpG conformational cycle ^55^, however further work is needed to validate this.

Collectively, this work demonstrates that Hsp90 provides a mechanism by which clients can escape the Hsp70-mediated folding block without compromising the ability of Hsp70 to prevent misfolding, in a process primarily driven by ATP hydrolysis. For HtpG, ATP binding alone is sufficient to drive the conformational rearrangements that prime it for ATP hydrolysis ^14,55^. However, in eukaryotes the conformational rearrangements and kinetics of ATP hydrolysis are exquisitely regulated by a complex host of co-chaperones ^56,57^, which allows the timing and outcome of Hsp90 activity to be fine-tuned according to the diverse folding requirements of individual clients. Despite this additional complexity, our findings indicate that ATP hydrolysis of Hsp90 remains the core conserved mechanism underlying client remodeling and as such, it is expected that the mechanistic insights provided here are evolutionarily conserved.

## Acknowledgments

We would like to thank the staff at Molecular Horizons for their technical and administrative support. This work was funded by the Australian Research Council (DP220103466). We also thank Prof. Matthias Mayer (University of Heidelberg Germany) for the provision of chaperone plasmid constructs and Dr. Lisanne Spenkelink (University of Wollongong, Australia) for her insightful feedback and suggestions on the manuscript. AMvO and HE also acknowledge funding from the National Health and Medical Research Council (APP1197069, APP1194872). D.C. was supported by the Lady Edith Wolfson Junior Non-Clinical Research Fellowship awarded by the MND Association UK (Cox 971-799), and Australian Research Council Discovery Early Career Researcher Award (DE240100707).

## Author contributions

**NRM**: Conceptualization (lead); Methodology (lead); Investigation (lead); Data curation (lead); Formal analysis (lead); Visualization (lead); Writing – original draft (lead); Writing – review and editing (equal). **BS**: Investigation (supporting); Conceptualization (supporting); Writing – review and editing (equal). **SM**: Investigation (supporting); Writing – review and editing (equal). **LR**: Formal analysis (supporting); Writing – review and editing (equal). **DC**: Formal analysis (supporting); Writing – review and editing (equal). **AMvO**: Conceptualization (supporting); Writing – review and editing (equal). Funding acquisition (equal). **HE**: Conceptualization (supporting); Writing – review and editing (equal). Funding acquisition (equal).

