## Supplementary files for "Direct single-molecule visualization of Hsp90-mediated relief of a Hsp70-folding block"

**Author notes**

* Lead contacts

**Contact information**

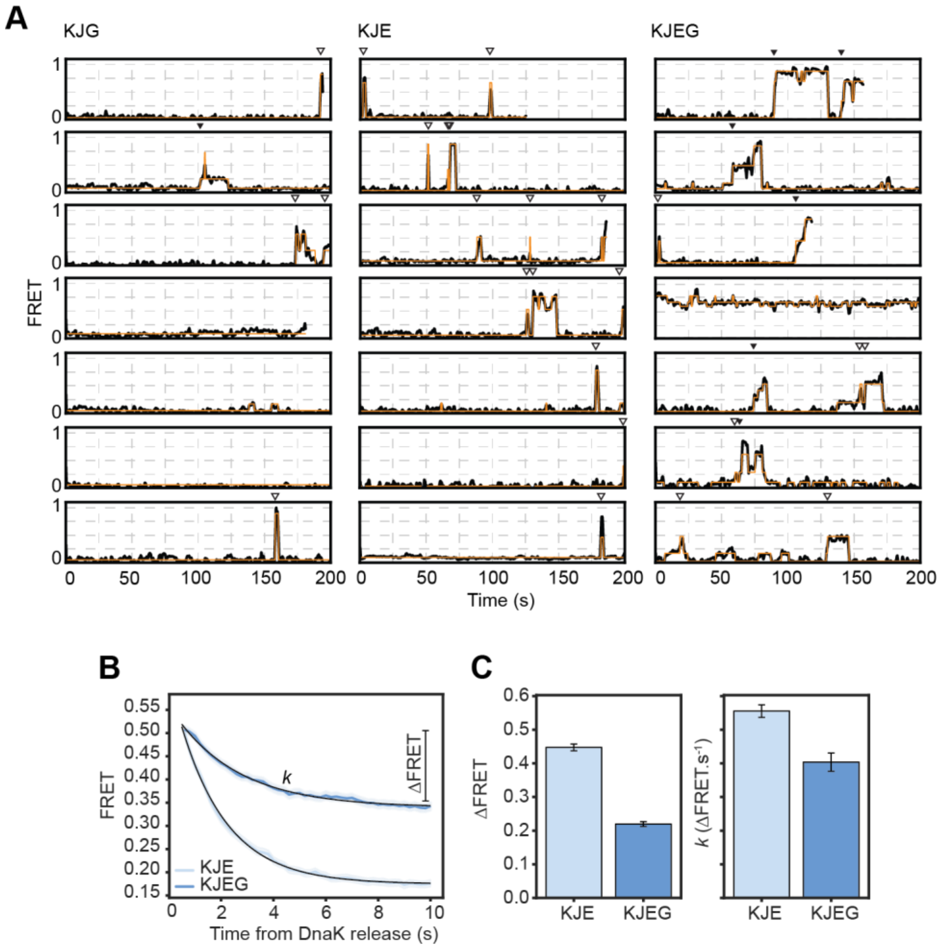


**Fig S1: HtpG reduces the rate of DnaK-rebinding and promotes efficient client refolding following DnaK release. (A)** Example FRET trajectories from individual $\text{Fluc}^{IDS1}$ molecules upon incubation with the indicated combination of molecular chaperones (HMM shown in orange). Non-controlled (*open triangles*) or controlled (*closed triangles*) DnaK-release events are shown above the traces. **(B)** Exponential fit of the FRET efficiency immediately following non-controlled DnaK-release from Fig 1G. **(C)** Both the change in FRET (*left*) and rate constant *k* (*right*) determined from the exponential fits in panel C are shown. Data is presented as mean ± standard deviation of fit.

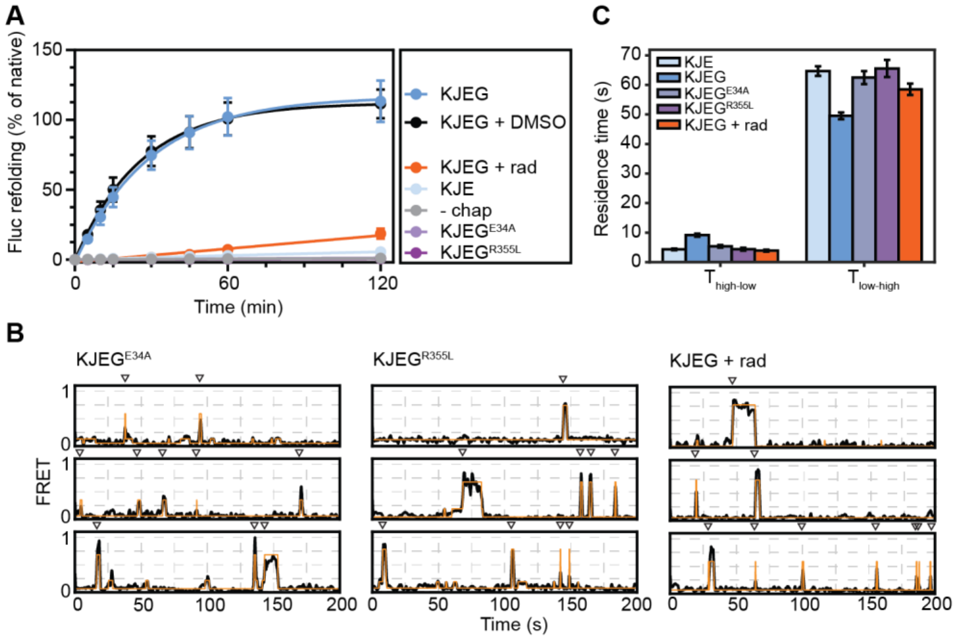


**Fig S2: HtpG interacts with DnaK and hydrolyzes ATP during productive client refolding. (A)** Luciferase refolding assay in the absence or presence of the KJEG system containing the indicated HtpG mutants or the HtpG inhibitor, radicicol (60 µM). Fluc$^{IDS1}$ was diluted 100-fold into refolding buffer alone (i.e., No chap) or supplemented with various combinations of molecular chaperones. All treatments contained 5 mM ATP. Data shown represents the mean ± SEM from three independent experiments. **(B)** Example FRET trajectories from individual Fluc$^{IDS1}$ molecules upon incubation with the indicated combination of molecular chaperones (HMM shown in orange). **(C)** Residence time data showing the time Fluc$^{IDS1}$ remains in a DnaK-bound state (i.e., < 0.3 FRET) prior to a non-bound state (i.e., $T_{low\text{-}high}$) and vice-versa. Data is presented as mean ± SEM from at least three independent experiments.

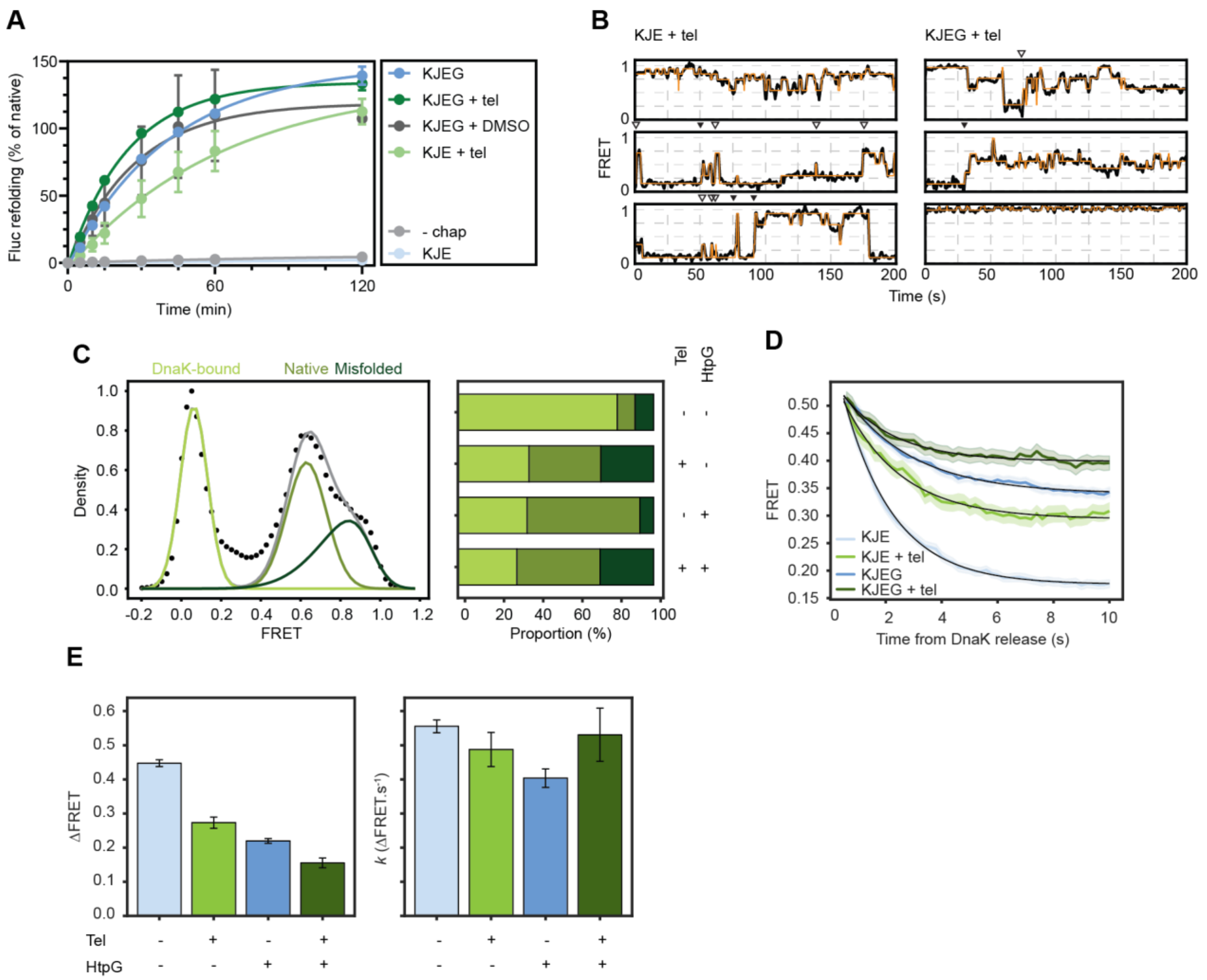


**Fig S3: HtpG mimics the telaprevir-induced inhibition of DnaK to enable client folding but does not prevent DnaK rebinding to misfolded proteins. (A)** Luciferase refolding assay in the absence or presence of the KJE or KJEG system that has been previously incubated with or without the DnaK-inhibitor telaprevir (100 µM). $Fluc^{IDS1}$ was diluted 100-fold into refolding buffer alone (i.e., no chap) or supplemented with various combinations of molecular chaperones. All treatments contained 5 mM ATP. Data shown represents the mean ± SEM from three independent experiments. **(B)** Example FRET trajectories from individual $Fluc^{IDS1}$ molecules upon incubation with the indicated combination of molecular chaperones and the DnaK-inhibitor telaprevir (100 µM, HMM shown in orange). **(C)** An example FRET efficiency KDE distribution dataset with the multiple-Gaussian model fits used to determine the proportion of $Fluc^{IDS1}$ states (i.e., native, misfolded or DnaK-bound, *left*) during chaperone-assisted refolding. The proportion of each state is shown (*right*) in the presence of the KJE system with or without HtpG and/or telaprevir. **(D)** Exponential fit of FRET efficiency immediately following non-controlled DnaK-release from Fig 2H. **(E)** Both the change in FRET (*left*) and rate constant *k* (*right*) as determined from the exponential fits in panel C are shown. Data for $Fluc^{IDS1}$ in the absence or presence of HtpG (i.e., KJE or KJEG) without telaprevir is the same as presented in Fig 1.

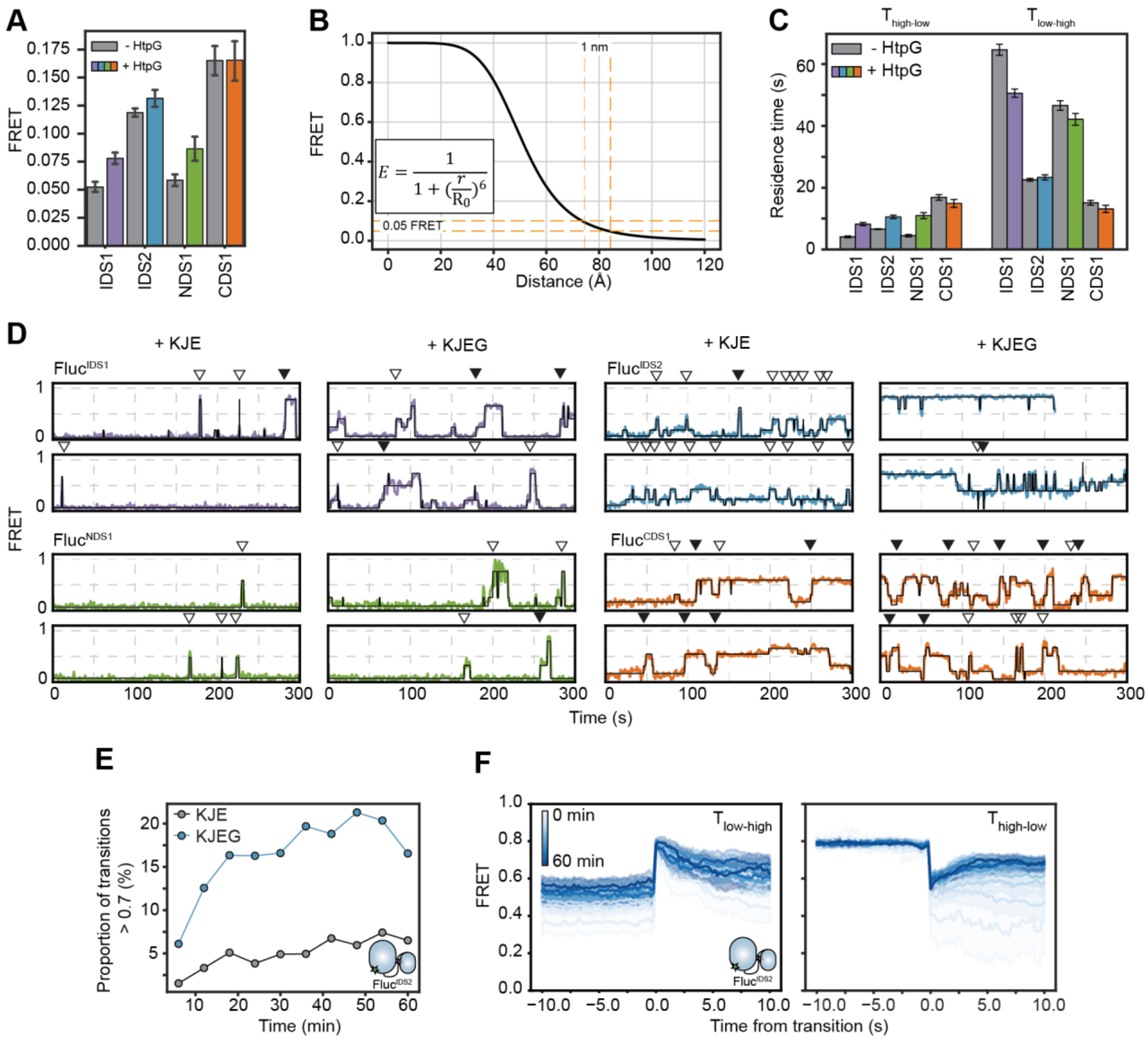


**Fig S4: Kinetics of HtpG-mediated folding of multiple Fluc domains. (A)** FRET efficiency of each Fluc sensor in the absence or presence of HtpG prior to DnaK-release events. **(B)** Non-linear dependence of FRET efficiency with distance. The FRET curve assumes a Forster radius of 51 Å (equivalent to that of an AF555/AF647 FRET-pair). Dotted lines demonstrate that the small change in FRET observed in panel A reports on a large difference in distance. The FRET equation is also shown, where *r* and $R_0$ are the distance between fluorophores and the Forster radius, respectively. **(C)** Residence time data showing the time each Fluc sensor remains in a DnaK-bound state (i.e., < 0.3 FRET) prior to a non-bound state (i.e., $T_{low\text{-}high}$) and vice-versa in the absence or presence of HtpG. Data is presented as mean ± SEM. **(D)** FRET trajectories for each Fluc sensor when incubated in the presence of the KJE system only or when supplemented with HtpG. **(E)** The proportion of transitions to the native $Fluc^{IDS2}$ state (> 0.7 FRET) in the absence or presence of HtpG over time. **(F)** The average FRET efficiency of all transitions towards (*left,* $T_{low\text{-}high}$) or away from (*right,* $T_{high\text{-}low}$) the native $Fluc^{IDS2}$ state (> 0.7 FRET) during refolding.

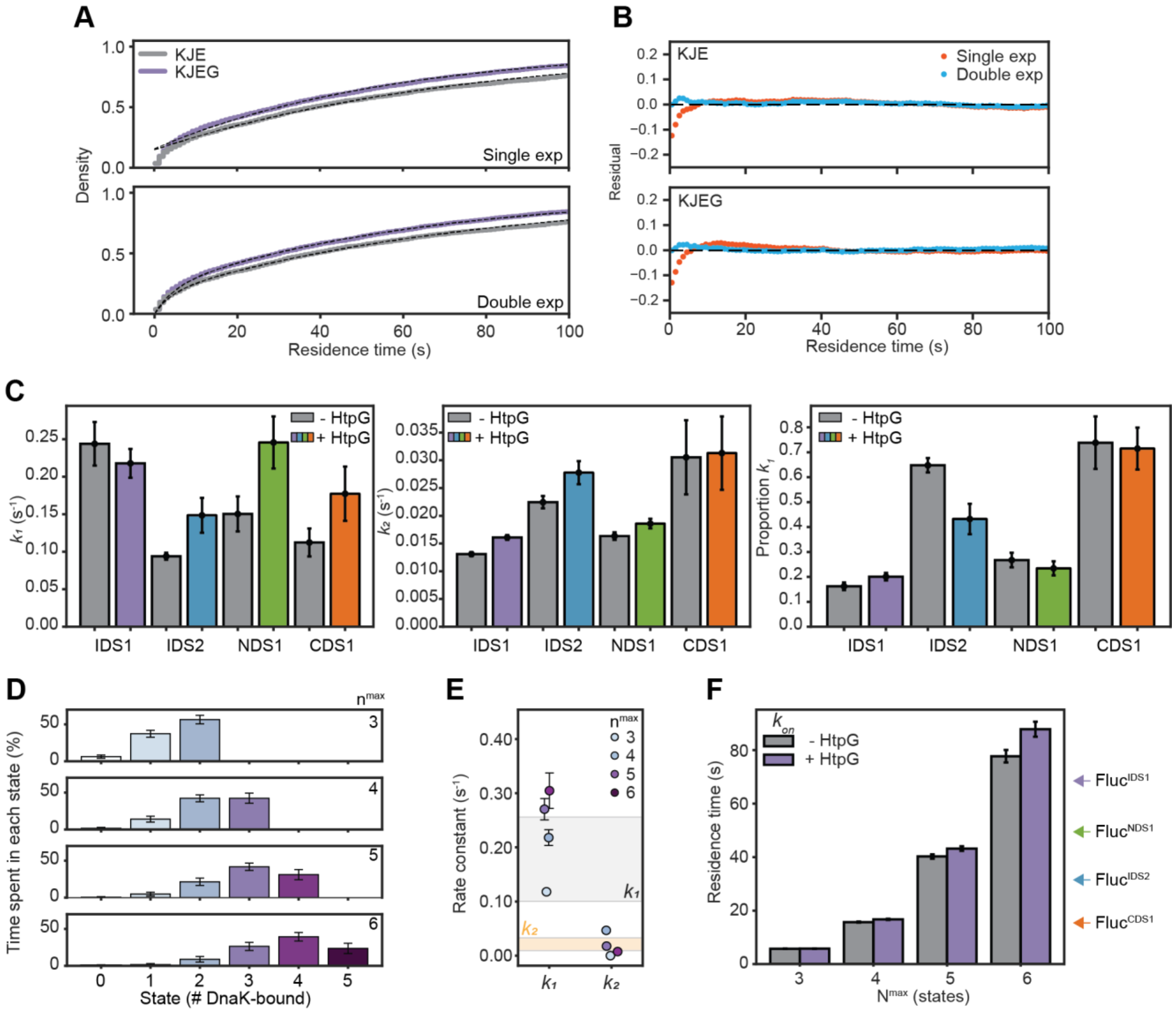


**Fig S5: Conformational compaction of Fluc upon DnaK-release is described by two rate constants. (A)** Cumulative density of Fluc$^{IDS1}$ $T_{low\text{-}high}$ residence times (from Fig S3D) in the absence or presence of HtpG, fit to either a (*top*) single- or (*bottom*) double-exponential model. **(B)** The residuals for both the single- and double-exponential fits from panel A are shown. **(C)** The extracted rate constants from the double-exponential fits for all Fluc sensors in the absence or presence of HtpG, with $k_1$ (*left*) and $k_2$ (*middle*) denoted. The proportion of the total fit that is described by $k_1$ is also shown (*right*). **(D)** The percentage of time that Fluc resides in each DnaK-bound state as a function of $state_{max}$, based on > 500 trace simulations. Data represents the mean ± standard deviation. **(E)** Predicted $k_1$ and $k_2$ rate constants following a double-exponential fit of $T_{low\text{-}high}$ residence times from simulated FRET trajectories at different $state_{max}$. Experimental rates for $k_1$ and $k_2$ determined from smFRET experiments are shown for comparison. Data represents the mean ± SEM. **(F)** The DnaK association rate (i.e., $k_{on}$) was determined in the absence or presence of HtpG (from Fig S1B-C) and used to simulate transitions between different DnaK-bound states and theoretical FRET trajectories. Simulations were performed with increasing $state_{max}$ and the mean $T_{low\text{-}high}$ residence times ± SEM from simulated FRET trajectories are shown. Mean $T_{low\text{-}high}$ residence times from smFRET experiments for each Fluc sensor in the absence of HtpG are shown for comparison.
